# Parallel genomic architecture underlies repeated sexual signal divergence in Hawaiian *Laupala* crickets

**DOI:** 10.1101/732966

**Authors:** Thomas Blankers, Kevin P. Oh, Kerry L. Shaw

## Abstract

When the same phenotype evolves repeatedly, we can explore the predictability of genetic changes underlying phenotypic evolution. Theory suggests that genetic parallelism is less likely when phenotypic changes are governed by many small-effect loci compared to few of major effect, because different combinations of genetic changes can result in the same quantitative outcome. However, some genetic trajectories might be favored over others, making a shared genetic basis to repeated polygenic evolution more likely. To examine this, we studied the genetics of parallel male mating song evolution in the Hawaiian cricket *Laupala*. We compared quantitative trait loci (QTL) underlying song divergence in three species pairs varying in phenotypic distance. We tested whether replicated song divergence between species involves the same QTL and the likelihood that sharing QTL is related to phenotypic effect sizes. Contrary to theoretical predictions, we find substantial parallelism in polygenic genetic architectures underlying repeated song divergence. QTL overlapped more than expected based on simulated QTL analyses. Interestingly, QTL effect size did not predict QTL sharing, but did correlate with magnitude of phenotypic divergence. We highlight potential mechanisms driving these constraints on cricket song evolution and discuss a scenario that consolidates empirical quantitative genetic observations with micro-mutational theory.

## INTRODUCTION

Parallel phenotypic evolution offers a unique opportunity to explore the possible routes from genotype to phenotype and variation in fitness among these routes. Common genetic mechanisms involved in parallel phenotypic evolution offer strong support for evolution by natural selection (Schluter and Nagel 1995; Losos 2011), because it is unlikely that stochastic processes such as mutation and drift would repeatedly act on the same genes. The extent of common genetic mechanisms also allows us to assess the predictability of evolutionary changes more broadly. Are the same genomic loci, genes, alleles, or nucleotides involved in the repeated evolution of the same trait? The answers to these questions are fundamentally important to evolutionary genetics and have wide-ranging applications, including in medicine (Nesse et al. 2012) and biodiversity management (Templeton 2017).

Current theory and evidence suggest that parallel phenotypic evolution, i.e. repeated divergence of the same phenotype in independent species pairs [see also ref (Conte et al. 2015)], can be explained by shared (“parallel”) genetic mechanisms (Conte et al. 2012; Martin and Orgogozo 2013). Theoretical work has shown that the probability of parallel genetic evolution from both *de novo* mutations and standing genetic variation increases with increasing strength of selection, increasing allelic effect sizes, decreasing numbers of loci, and higher starting allele frequencies (Orr 2005b; Chevin et al. 2010; MacPherson and Nuismer 2017). Accordingly, several traits that experience strong ecological selection pressures and are controlled by one or few genetic loci have been found to evolve repeatedly by changes in the same gene, e.g. armor plating in stickleback by *eda* (Colosimo et al. 2005) or wing pattern mimicry in butterflies by *optix* (Reed et al. 2011). Conversely, parallel phenotypic evolution of traits controlled by many loci of small effect and mostly evolving through soft rather than hard selective sweeps (Barrett and Schluter 2008; Messer and Petrov 2013), should be less likely to have a shared genetic basis. Accordingly, several empirical studies have supported the hypothesis of an independent genetic basis for polygenic traits (Deagle et al. 2011; Renaut et al. 2011; Kautt et al. 2012; Westram et al. 2014; Ravinet et al. 2016). Likewise, simulations have shown that independent polygenic divergence events draw on different combinations of alleles to achieve parallel phenotypic outcomes (Yeaman and Whitlock 2011), resulting in limited shared variants.

However, some empirical support for parallel polygenic genetic architectures exists, ranging from reuse of QTL (Conte et al. 2015), to genes (Tenaillon et al. 2012; Kryazhimskiy et al. 2014; Yeaman et al. 2016), and even nucleotides (Tennessen and Akey 2011; Graves et al. 2017). One reason to expect parallel genetic architectures for repeated polygenic evolution is that high levels of shared ancestral polymorphisms, characteristic of rapid species radiations, can be recruited repeatedly in parallel divergence events, especially when certain genetic combinations carry additional fitness penalties due to pleiotropy and linkage (Elmer and Meyer 2011; Conte et al. 2012; Marques et al. 2018). Moreover, during divergence, especially when some gene flow occurs between species or populations, the migration-selection-drift balance may consolidate quantitative and otherwise polygenic phenotypic variation into relatively few QTL of large effect (Yeaman and Whitlock 2011). This happens because even if small-effect loci initially contribute to variation, these tend to be replaced by loci of larger effect, which are more favorable since there are fewer cross-over events that can result in maladaptive combinations. Genomic rearrangements may further consolidate co-adaptive alleles, resulting in small effect alleles clustered into tightly linked groups of loci that effectively inherit as a single QTL of large effect (Yeaman and Whitlock 2011; Yeaman 2013). Thus, even if there are many possible genetic trajectories leading to phenotypic change, these trajectories may have differential fitness, the outcome being that evolution proceeds along the lines of least resistance (Schluter 1996). However, empirical studies of the mechanisms and genetic architecture underlying replicated polygenic divergence are rare in the empirical literature, even though much of life’s complexity is quantitative and evolves by many genetic changes of small effect [*i.e.* polygenic genetic architectures; (Mackay 2001; Orr 2005a)]. This is especially true for traits involved in divergent sexual behaviors, despite their prominent role in repeatedly driving natural variation across the animal kingdom (Ritchie and Phillips 1998; Coyne and Orr 2004).

Here, we examine the extent of parallel genetic differentiation underlying mating song divergence among phylogenetically independent pairs of species (independent contrasts) of the Hawaiian swordtail cricket genus *Laupala* (Fig 1A). Species of the genus *Laupala* are morphologically cryptic, flightless, forest-dwelling crickets with geographic ranges endemic to single islands within the archipelago. The male mating songs of these crickets are simple pulse trains produced by single wing strokes at regular intervals (Otte 1994). The diversification of male mating song in *Laupala* is quantitative in nature, largely due to pulse rates ranging from relatively slow to relatively fast. Moreover, the songs of closely related species pairs, as well as species in sympatry, always differ (i.e. are “slow” or “fast” relative to each other). With the results of two new QTL experiments, and integrating results from two previous studies (Shaw et al. 2007; Blankers et al. 2018b), we compare the location and effect of pulse rate QTL across three independently diverged species pairs (six species in total). We test two predictions emerging from the theoretical and empirical work discussed above: (1) when the potential for numerous, small genetic changes underlies phenotypic evolution, as is expected for complex traits, repeated divergence events in such traits are unlikely to share a common genetic basis (e.g. (Elmer and Meyer 2011; Agrawal 2017)); and (2) when loci are shared among divergence events, they will be those of relatively larger phenotypic effect (Orr 2005b; MacPherson and Nuismer 2017).

**Figure 1.**
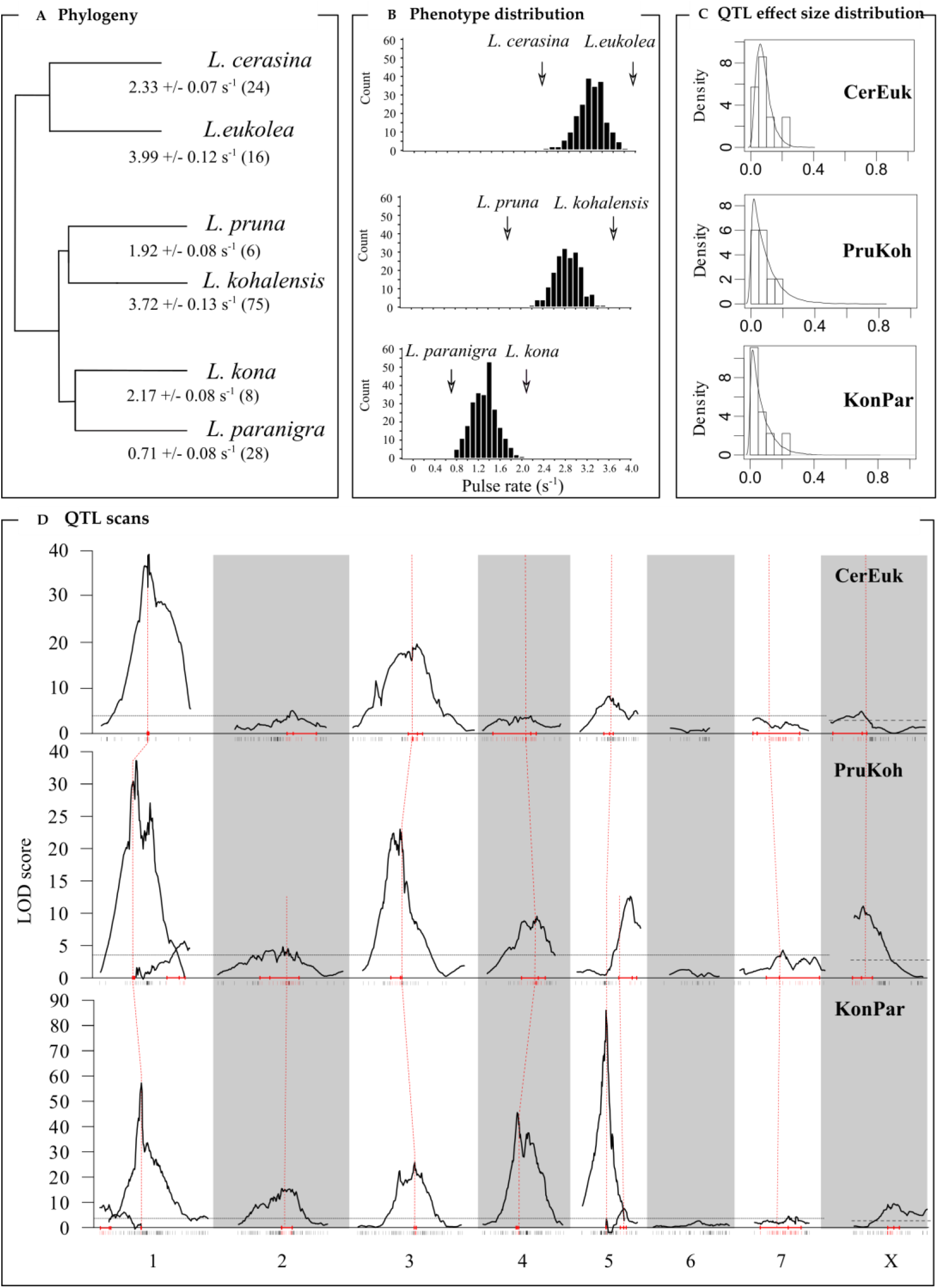
Phenotypic divergence and QTL scan. (A) Phylogenetic relationships [based on (Mendelson and Shaw 2005)] for the three species pairs in this study along with species averages and standard deviations for the male pulse rate. Sample sizes for song recordings are in parentheses. (B) Phenotypic distribution of the F_2_ progeny used for QTL mapping. Data sources: CerEuk [top, from ref (Blankers et al. 2018b)], PruKoh (middle; this study), and KonPar (bottom; this study). Arrows indicate the average phenotypic value of the parental lines of the intercross families. (C) Probability density functions (solid lines) of the best fit exponential (KonPar, PruKoh) and gamma distribution (CerEuk) and the density histogram (bars) for the effect size distribution. (D) QTL scan. The strength of linkage (LOD score) of the markers with pulse rate variation across the eight linkage groups is shown for the multiple QTL model (black lines). Horizontal lines indicate the significance threshold at 5% FDR. Red horizontal lines on the x-axis show the location of the QTL peak (center vertical bar) and the limits of the 95% BCI interval. Red dashed lines connect markers close to or at the QTL peak that are shared between one or more crosses (see also Table S2, S3, and Fig S1).

## METHODS

### Samples

Comparative QTL mapping was done combining results from a previously published QTL study involving an interspecies cross between *Laupala cerasina* and *L. eukolea* (hereafter referred to as “CerEuk”) (Blankers et al. 2018b), and from two new QTL analyses involving *L. kona* and *L. paranigra* (“KonPar”), and, *L. pruna* and *L. kohalensis* (“PruKoh”). All species diverged recently [less than 0.5 million years ago (Mendelson and Shaw 2005)] on the Hawai’i Island, except *L. eukolea*, which is an endemic on the neighboring island Maui. A combination of wild-caught and lab-reared individuals was used for the parental crosses. For the new analyses presented here, we crossed one male (*L. kona* or *L. pruna*) and one female (*L. paranigra* or *L. kohalensis*) of each species pair, then individually sib-mated multiple male and female first-generation hybrid offspring to generate 263 (KonPar) and 193 (PruKoh) second-generation hybrid (F2) males. Details about the other crosses are in references (Blankers et al. 2018a) and (Blankers et al. 2018b).

### Genotypes

DNA from parental and all second-generation hybrid offspring (KonPar: N = 263; PruKoh: N = 193; CerEuk: N = 230) was isolated using the DNeasy Blood & Tissue Kit (Qiagen, Valencia, CA, USA) and sequenced following the Genotype-By-Sequencing (GBS) protocol (Elshire et al. 2011) with *Pst* I, obtaining 100 bp reads on the Illumina HiSeq 2000 (Illumina, San Diego, CA). After quality control and processing of the raw reads, genotypes were called using an approach combining FreeBayes (Garrison and Marth 2012) and the Genome Analysis Toolkit (DePristo et al. 2011; Van der Auwera et al. 2013) after which genotypes were filtered based on stringent criteria and only retaining bi-allelic SNPs with less than 10% missing data. Further details for genotype calling and filtering have been described previously (Blankers et al. 2018b,a).

### Phenotypes

Songs were recorded in a temperature controlled, anechoic chamber (for KonPar, temperature range: 19.9 – 20.7° C; mean + standard deviation: 20.5 + 0.15 °C; for PruKoh, temperature range: 19.3 – 20.8° C; mean + standard deviation: 20.2 + 0.34 °C) with a Sony Pro cassette recorder and condenser microphone from screen/plastic chambers. The pulse period (the beginning of one pulse to the beginning of the following pulse) was measured from five independent periods (accurate to 10 ms) and transformed to pulse rates (the inverse of pulse period). Recording and quantification of song variation followed established procedures (Shaw 1996).

### Linkage maps

The linkage maps and detailed methods regarding their assembly have been described previously (Blankers et al. 2018a,b). In short, marker loci (ancestry informative loci not deviating significantly from 1:2:1 or 1:1 ratios for autosomal or X-linked inheritance, respectively) were grouped into linkage groups using JoinMap 4.0 (van Ooijen 2006) and MapMaker 3.0 (Lander et al. 1987; Lincoln et al. 1993) at LOD > 5. Then, for each linkage group, we used high confidence markers that had identical order in JoinMap and MapMaker to create a high confidence scaffolding map. The scaffolding map was then filled-out in MapMaker by iteratively adding markers at log-likelihood thresholds of 3.0, 2.0, and then 1.0, randomly permutating marker orders in 7 marker windows to explore alternative orders between each round.

### QTL scan

The first goal of this study was to find the number and location of genomic loci underlying pulse rate variation and estimate their contribution to phenotypic divergence. The QTL scans and comparative QTL analyses were done in R (R Development Core Team 2016) and custom codes are available at github.com/thomasblankers/QTLreuse. We detected QTL using single-QTL scans followed by multiple QTL mapping (MQM) in r/QTL as described previously (Blankers et al. 2018b). Thresholds for including additional QTL at false positive error rate lower than 0.05 were based on 1,000 permutations. QTL effect sizes were calculated from the MQM models using drop-one term ANOVA. Based on the total detected QTL and their effect sizes estimated from the MQM models, we estimated the true number of loci using the equations in (Otto and Jones 2000).

### QTL overlap

We tested whether any of the detected QTL in the three independent contrasts (CerEuk, KonPar, and PruKoh) overlapped. For each of the detected QTL, we calculated 95% Bayesian Credible Intervals. QTL reuse was inferred when BCIs overlapped between any two (referred to as ‘double QTL’ from hereon) or all three (‘triple QTL’) species pairs. QTL present in one cross that did not overlap with any other QTL were dubbed ‘unique QTL’. To test whether the observed extent of QTL overlap would be expected by chance, we first approximated the probability that we would find six overlapping QTL by chance, given the average size of a confidence interval, the average linkage group length, and the number of detected QTL. The probability was derived from 100,000 simulations where at each iteration two confidence intervals of *s* cM (average confidence interval size) were placed on a LG of length *l* cM (average linkage group length), *n* times (where *n* is the number of detected QTL) using custom code (https://github.com/thomasblankers/qtlreuse).

To more realistically reflect the conditions of our comparative QTL experiment, we also simulated the entire analysis (linkage mapping, QTL mapping, QTL overlap) 1,000 times and retained quantiles on the extent of QTL overlap when QTL effects are randomly distributed across the genome. We first obtained a simulated linkage map with 8 linkage groups (one of which was X-linked) of lengths varying between 150 cM and 75 cM and randomly placed markers on that map averaging a marker every 2 cM. We then randomly distribute 16 “true” loci with their effect sizes drawn from an exponential distribution with mean 0.047 pulses per second (corresponding to the overall mean effect size across all QTL detected in PruKoh, KonPar, and CerEuk) on that map. We simulated a QTL cross object using the simulated map and phenotypes for 200 F2 individuals. Using ‘stepwiseqtl’, we performed an automated version of multiple QTL mapping to identify and localize the randomly drawn QTL effects and calculated BCIs for each of the identified QTL. Through 1,000 iterations of two simulated QTL scans per iteration we tested if any one or more of the BCIs in the two simulated QTL scans overlapped and counted the number of overlapping BCIs in each iteration. Any BCI overlap here is purely stochastic as the effect size and location of the QTL in the two parallel simulated crosses are randomly drawn from a hypothetical distribution of QTL. However, the linkage map, the number of “true” loci, and the effect size distribution are based on our observations for the three crosses studied here (see Results).

### Parallel QTL effects

There is considerable variation in the magnitude of phenotypic divergence between parental lines across the three species crosses analyzed here, as well as a fourth cross, between *L. kohalensis* and *L. paranigra*, examined in a previous study (Shaw et al. 2007). The latter cross is not independent of the KonPar and PruKoh crosses but provides an interesting contrast with respect to increasing time since divergence and phenotypic distance (Fig 1A). The phenotypic difference between *L. kohalensis* and *L. paranigra* is 3.01 pulses per second, whereas parental lines of KonPar, CerEuk, and PruKoh differ by 1.46, 1.66, and 1.64 pulses per second, respectively. We asked whether the magnitude of phenotypic divergence was related to the (detected or estimated total) number of QTL, which is expected if pulse rate evolution occurs by accumulating variants in additional genomic regions, or by increasing the average QTL effect size, which is expected if pulse rate evolution occurs by accumulating variants in genomic regions where other causal variants already reside or by selection favoring variants of increasingly larger phenotypic effect in those regions.

To further examine the genetic basis of repeated pulse rate evolution, we examined effect size variation in shared and unique QTL. If QTL that are shared between species pairs contain a single causal variant that they have in common, the effects of those QTL are expected to be of similar magnitude in the two crosses. Using linear regression, we tested for significant correlations of effect size among shared QTL among species pairs. We regressed the vector with all effect sizes in one cross onto the vector of a second cross and asked whether QTL effect sizes were significantly correlated. Lastly, we tested whether QTL of larger effect are more likely to be shared. This prediction follows from theoretical research that suggests genetic parallelism is more likely when phenotypic effects are larger (Orr 2005b; Chevin et al. 2010; MacPherson and Nuismer 2017). We tested for a correlation between QTL effect size (calculated above) and the extent of sharing (‘unique’, ‘double’, or ‘triple’), while including species pair (‘KonPar’, ‘PruKoh’, ‘CerEuk’) as a covariate to account for species-specific effects.

## RESULTS

### Sequencing and linkage mapping

For the linkage maps, 650 (KonPar) and 325 (PruKoh) SNP markers were grouped into eight linkage groups at a log-of-odds (LOD) threshold of 5.0. Using a combination of Joinmap (van Ooijen 2006) and MapMaker (Lander et al. 1987; Lincoln et al. 1993) to iteratively order markers within linkage groups at decreasing stringency of linkage criteria, we obtained two collinear maps (Fig S1). The total map lengths were 887.3 cM (KonPar) and 1056.3 cM (PruKoh), corresponding to an average marker spacing of 1.37 and 3.25 markers/cM, respectively (Fig S1). The average sequencing depth (± standard deviation) per linkage map marker across individuals was 38.1x (± 23.8), and 41.8x (± 29.3) for KonPar and PruKoh, respectively. More detailed results on the linkage maps (and collinearity) have been published previously (Blankers et al. 2018a).

### Is divergence in replicate species pairs associated with similar genetic architectures?

In line with previous results for pulse rate variation in *Laupala*, our findings support a polygenic, additive genetic architecture. Second-generation lab-reared hybrids have phenotypic values (i.e. pulse rates) that are intermediate relative to the parental lines (Fig 1B). Previously, we detected seven QTL for CerEuk (N = 230; Fig 1B top panel, Table S1). Our current QTL analyses revealed eight QTL for PruKoh and nine QTL for KonPar and effect size distributions follow either an exponential or gamma distribution (Fig 1C,D; Table 2). In addition, two significant interactions were found for KonPar (but not for any of the other crosses), involving QTL on LG1 and LG4, and on LG1 and LG5 (Table 1). In all three crosses, multiple loci of small (< 5% of the parental difference) to moderate (< 17% of the parental difference) effect were found on the same 6 autosomal (all but LG6) and the X-linked linkage groups (Table 1, Table 2, Table S1).

**Table 1.**
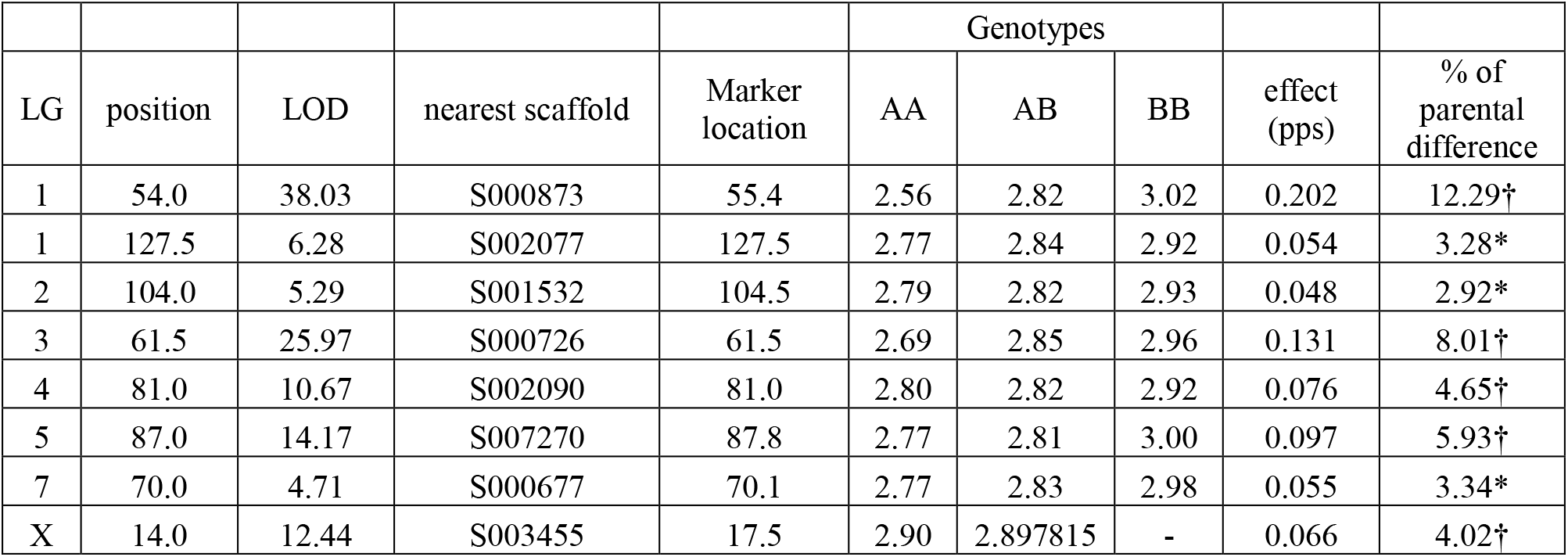
QTL results for PruKoh. QTL were mapped using 193 F2 individuals. LG: linkage group. LOD: log-of-odds. A and B alleles denote *L. pruna* and *L.kohalensis* alleles, respectively. QTL effects are shown as the estimated effect in pulses per second derived from the multiple QTL model and as the percent variance explained relative to the parental difference (1.64 pps). All QTL effects are significantly different from zero (* P < 0.05; † P < 0.01).

|  |  |  |  |  | Genotypes |  |  |  |  |
| --- | --- | --- | --- | --- | --- | --- | --- | --- | --- |
| LG | position | LOD | nearest scaffold | Marker location | AA | AB | BB | effect (pps) | % of parental difference |
| 1 | 54.0 | 38.03 | S000873 | 55.4 | 2.56 | 2.82 | 3.02 | 0.202 | 12.29† |
| 1 | 127.5 | 6.28 | S002077 | 127.5 | 2.77 | 2.84 | 2.92 | 0.054 | 3.28* |
| 2 | 104.0 | 5.29 | S001532 | 104.5 | 2.79 | 2.82 | 2.93 | 0.048 | 2.92* |
| 3 | 61.5 | 25.97 | S000726 | 61.5 | 2.69 | 2.85 | 2.96 | 0.131 | 8.01† |
| 4 | 81.0 | 10.67 | S002090 | 81.0 | 2.80 | 2.82 | 2.92 | 0.076 | 4.65† |
| 5 | 87.0 | 14.17 | S007270 | 87.8 | 2.77 | 2.81 | 3.00 | 0.097 | 5.93† |
| 7 | 70.0 | 4.71 | S000677 | 70.1 | 2.77 | 2.83 | 2.98 | 0.055 | 3.34* |
| X | 14.0 | 12.44 | S003455 | 17.5 | 2.90 | 2.897815 | - | 0.066 | 4.02† |

**Table 2.** QTL results for KonPar. QTL were mapped using 263 F2 individuals. LG: linkage group. LOD: log-of-odds. A and B alleles denote *L. paranigra* and *L.kona* alleles, respectively. QTL effects are shown as the estimated effect in pulses per second derived from the multiple QTL model and as the percent variance explained relative to the parental difference (1.46 pps). All QTL effects are significantly different from zero (* P < 0.05; † P < 0.01) except for the LG5 x LG1 interaction (which increases overall QTL model fit, but the slope the relationship between genotype interaction and phenotypic effect size is not significant).

| LG | position | LOD | nearest scaffold | Marker location | Genotypes |  |  | effect (pps) | % of parental difference |
| --- | --- | --- | --- | --- | --- | --- | --- | --- | --- |
|  |  |  |  |  | AA | AB | BB |  |  |
| 1 | 12.7 | 9.65 | S002179 | 12.7 | 1.25 | 1.33 | 1.34 | 0.035 | 2.39* |
| 1 | 58.1 | 62.88 | S002151 | 58.1 | 1.17 | 1.33 | 1.44 | 0.112 | 7.64† |
| 2 | 60.0 | 16.70 | S000517 | 60.1 | 1.27 | 1.30 | 1.39 | 0.048 | 3.26† |
| 3 | 79.1 | 28.08 | S001173 | 79.1 | 1.27 | 1.29 | 1.41 | 0.070 | 4.80† |
| 4 | 48.0 | 49.93 | S001965 | 48.8 | 1.22 | 1.28 | 1.47 | 0.096 | 6.57† |
| 5 | 32.5 | 94.61 | S004383 | 32.5 | 1.07 | 1.34 | 1.55 | 0.217 | 14.88† |
| 5 | 57.6 | 8.06 | S000770 | 57.6 | 1.18 | 1.31 | 1.49 | 0.019 | 1.29† |
| 7 | 47.0 | 4.65 | S001185 | 46.9 | 1.29 | 1.33 | 1.31 | 0.023 | 1.58* |
| X | 55.0 | 10.11 | S00105 | 53.4 | 1.36 | 1.27 | 1.36 | 0.025 | 1.72* |
| LG5 x LG1 | (interaction) | 6.659 | - | - | - | - | - | 0.002 | 0.13 |
| LG1 x LG4 | (interaction) | 10.042 | - | - | - | - | - | -0.025 | -1.71* |

Using the framework in (Otto and Jones 2000), we estimated the total number of QTL in each cross (assuming an exponential model and estimating the detection threshold, θ, for QTL from the data). The predicted total number of QTL was 12.05 (95% confidence interval: 5.79 – 21.72; θ = 0.0135) for KonPar and 19.78 (9.05 – 36.83; θ = 0.0418) for PruKoh. Based on the data in Table 3 in (Blankers et al. 2018b), the total number of QTL for CerEuk is estimated at 16.60 (7.14 – 32.12).

### Are the same QTL regions involved?

To account for the uncertainty associated with estimating QTL peak locations, we considered QTL to be shared between two (‘double’ QTL) or all three (‘triple’ QTL) species pairs if the 95% Bayesian Credible Interval (BCI) for QTL on the homologous linkage groups overlapped. Out of a total of 12 QTL detected in the three species crosses combined, we found four triple QTL and four double QTL (two between PruKoh and KonPar, one each between those and CerEuk) and four QTL unique to a single cross (Fig 1; Table S4; Fig S2 – S4). The probability of finding six overlapping confidence intervals of average size 21.8 cM for a total of 7 detected QTL that are each located on a separate linkage group of average size 121.5 cM is 0.004. If there are a total of 9 QTL, this probability increases to 0.025.

To account for the fact that there are more possible QTL than the ones we detect, we calculated the probability that the number of shared QTL (six between KonPar and PruKoh, five between CerEuk and both KonPar and PruKoh) would be observed by chance, given the true number of QTL (between 12 and 20) and the detected QTL (between seven and nine). Ninety-five percent of the simulated comparative QTL experiments resulted in fewer than four shared QTL and only 1% resulted in six shared QTL. In addition, we found that shared QTL in simulations were typically those with relatively broad confidence intervals, while in the data we observe overlap even between QTL with relatively narrow confidence intervals (e.g. LG1 and LG3; Fig 1C). Overall these results suggest that the level of QTL overlap observed here (four QTL shared between all three crosses and an additional one or two shared in pairwise comparisons) is higher than expected by chance.

Despite significant overlap in QTL locations, phenotypic effects of shared QTL varied among the crosses. We inspected the relationship between phenotypic effect size, the number of QTL, and the magnitude of phenotypic divergence between the parental species. The phenotypic difference was unrelated to the number of detected QTL (r^2^ = 0.01, F_1,2_ = 0.03, *P* = 0.8862) or the estimated true number of loci (r^2^ = 0.10, F_1,2_ = 0.23, *P* = 0.6797), but was significantly associated with the average effect size (in pulses per second) across all detected QTL (r^2^ = 0.96, F_1,2_ = 68.01, *P* = 0.0144; Fig 2A). Effect sizes were significantly correlated between QTL shared by PruKoh and CerEuk (r^2^ = 0.99, F_1.3_ = 326.4, P = 0.0004) but not in the other two comparisons (KonPar and PruKoh: r^2^ = 0.42, F_1.5_ = 2.92, P = 0.1626); CerEuk and KonPar: r^2^ = 0.004, F1.4 = 0.012, P = 0.9181; Fig 2B). Furthermore, we found no association between effect size and QTL sharing (partial R^2^ = 0.02, F_4,19_ = 0.83, P =0.5222; Fig 2C). *Post hoc* tests further revealed that average effect sizes did not differ significantly among any categories of QTL sharing. This indicates that QTL reuse is not biased towards QTL of larger effect.

**Figure 2.**
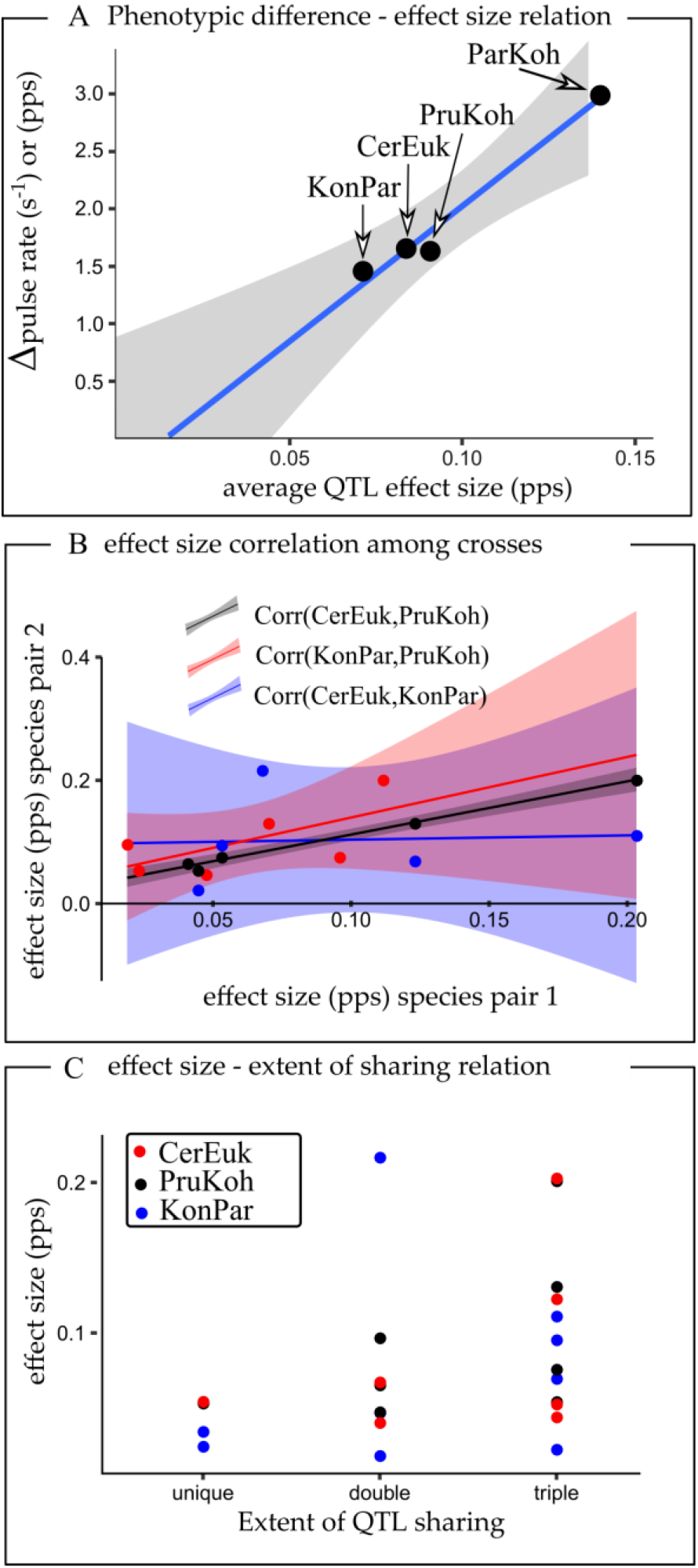
Shared QTL effects. (A) Association between phenotypic divergence (the difference between parental pulse rates) and the average effect size of detected QTL across (from lowest to highest) KonPar, CerEuk, PruKoh, and ParKoh. The latter data are from Table 3 in (Shaw et al. 2007). (B) For each of the pairwise comparisons between QTL crosses, the effect size of the shared QTL (circles), measured as the increase/decrease in pulse rate per alternative allele, is shown for both crosses. The solid line is a linear regression on these data. (C) The effect size of QTL (dots) across cross families (colors) is not related to the extent of QTL sharing. However, QTL not shared among any species pairs tend to be of smaller effect than those that are shared across two or all three species pairs (not significant).

## DISCUSSION

In the evolution of complex traits, multiple genetic trajectories can achieve the same net phenotypic effect. However, repeated use of the same genetic elements underlying parallel phenotypic evolution suggests that there are only few feasible genetic pathways towards phenotypic change. Such genetic constraints can arise due to limited molecular pathways by which specific evolutionary changes can be achieved, limited genetic variation, as well as variable fitness potential of the possible genetic trajectories (Elmer and Meyer 2011; Yeaman et al. 2018). Thus far, theoretical and empirical research primarily has focused on the recurrent use of major effect loci that are under strong ecological selection (Stern and Orgogozo 2008; Conte et al. 2012; Martin and Orgogozo 2013; Stern 2013). This focus leaves important gaps in our knowledge about genetic parallelism, particularly about traits controlled by many genes of small effect. Such traits are common in nature and central to quantitative genetic theory (Robertson 1967; Lande 1981; Mackay 2001), and include, for example, mating behavior and ecological adaptations that often characterize early phases of the speciation process (Coyne and Orr 2004).

Here we asked whether the genetic architecture underlying mating song divergence in Hawaiian swordtail crickets (*Laupala*) is shared among phylogenetically independent species pairs. The *Laupala* radiation is a powerful study system to address the genetics of parallel phenotypic evolution. The 38 morphologically and ecologically cryptic species, which display robust species boundaries in the wild but can interbreed in the lab, have evolved conspicuous differences in mating signals and preferences on short evolutionary timescales. Recent, repeated evolution in this significant barrier to gene flow provides a fascinating natural context to study the evolutionary genetics of sexual selection and speciation.

We find that repeated and independent interspecific divergences of *Laupala* mating songs involve broadly similar genetic architectures (Fig 1) and have more QTL in common than would be expected by chance. Therefore, we reject the hypothesis that repeated episodes of divergence in polygenic traits travel along unique evolutionary genetic trajectories, providing important evidence for genetic constraints on polygenic evolution of courtship song. Our result agrees with previous studies in this system in that pulse rate divergence in *Laupala* is caused by many, additive loci of small to moderate effect (Shaw et al. 2007; Ellison et al. 2011; Oh et al. 2012), but goes further to show common QTL underlying these changes. Our study further shows that effect size does not predict whether a QTL is likely to be reused in independent, parallel divergence events (Fig 2). Consequently, we also reject the hypothesis based on recent theoretical work (MacPherson and Nuismer 2017) that loci of relatively larger effect are more likely to diverge in parallel than those of smaller effect.

### Genomic constraints on polygenic evolution

Our results indicate genomic constraints to polygenic sexual signal evolution in *Laupala* crickets. First, we found that at least five QTL overlapped between any species pair. Our simulations suggest that the probability of observing five or six QTL in common between crosses is less than 5%, given the size of linkage groups and QTL confidence intervals, the number of QTL, and their effect size. These results thus provide evidence that QTL from overlapping regions of homologous linkage groups are recruited repeatedly as species diverged in pulse rate. With current data, while we *can* conclude that these changes occur within the same QTL regions, at least some of which span just a few cM, whether they have occurred in the same genes must await identification of the specific genes involved.

Second, comparing QTL effect sizes between species pairs sharing QTL revealed that when phenotypic divergence increases over time, this is achieved not by recruiting additional QTL, but rather by larger effects at existing QTL. Average effect size, across all QTL, scales linearly with the magnitude of the pulse rate difference (Fig 2A). Conversely, we found no relationship between the average QTL effect size and the number of detected QTL that explain species differences in pulse rate. Thus, elaboration of pulse rate differences is associated with larger effects of the same QTL, rather than with additional QTL elsewhere in the genome. Furthermore, phenotypic effects at shared QTL varied substantially between crosses and were uncorrelated (Fig 2B). Effect size variation among shared QTL suggests that these regions contain distinct, but closely linked, variants or combinations of variants. Because average QTL effect size in each cross and the associated total phenotypic difference increase with increasing divergence times, a reasonable hypothesis is that QTL regions harbor multiple, tightly-linked variants of small effect that have accumulated over time.

There are two leading candidate explanations for the observed molecular constraints. First, spatial limits on where in the genome variation is generated (e.g. through *de novo* mutations and recombination of standing genetic variation), maintained (e.g. through balancing selection), and selected (Manceau et al. 2010; Losos 2011; Martin and Orgogozo 2013; Stern 2013) may limit the genetic trajectories towards parallel phenotypic evolution. These constraints may arise as a consequence of mutation rate variation and epistatic and pleiotropic interactions in specific genomic regions that ultimately can strengthen the efficacy of selection (Kopp 2009; Manceau et al. 2010; Martin and Orgogozo 2013; Stern 2013). Moreover, genes controlling mating song rhythm in crickets may be spatially concentrated in functional genetic clusters, making some of the above genetic conditions more likely. Tight linkage of multiple genes underlying a polygenic phenotype would render inheritance of those genes non-independent, constraining the number of segregating genomic regions that can contribute to phenotypic variation. Parallel phenotypic change would result from the same or different substitutions in the same cluster. Interestingly, multiple substitutions may accumulate across genes within a cluster, and jointly contribute to the phenotypic effect of the QTL (Huxley 1942; Stern 2000), especially if genomic rearrangements are favorable under the selection-migration balance (Yeaman and Whitlock 2011; Yeaman 2013). There is evidence that song rhythm QTL in *Laupala* fall in regions enriched for genes with neuronal and motor rhythmic functions (Blankers et al. 2018b), suggesting that pulse rate differentiation might be controlled by clusters of closely linked genes.

Second, sexually selected phenotypes such as pulse rate in the crickets studied here (Oh and Shaw 2013) might be prone to genetic parallelism due to interactions with other genotypes or phenotypes. This follows because divergence in sexual phenotypes often requires trait-preference co-evolution for the system to remain functional during evolutionary change, i.e. coordinated divergence in signaling and receiver traits (Lande 1981; Kirkpatrick 1982). One way that coordinated change between signal and preference might be maintained during divergence is for substitutions and mutations to occur only in those signal genes that are linked to the preference genes. This would limit the number of possible fitness enhancing variants and thereby increase the likelihood of genetic parallelism. Interestingly, there is evidence that supports colocalization (i.e. tight linkage or pleiotropy) of pulse rate and pulse rate preference QTL in the *Laupala* genome (Shaw and Lesnick 2009; Wiley et al. 2012). It is thus conceivable that linkage or pleiotropy underlying the genes of male and female traits in *Laupala* are contributing to the patterns of shared QTL that we observed.

Another source of constraint on the molecular pathways towards phenotypic change is when adaptation results mostly from standing genetic variation, rather than from *de novo* variants. Our current data do not allow us to distinguish between the two alternatives. On the one hand, given the rapid evolution of pulse rates and relatively low intraspecific versus high interspecific variation in *Laupala* (see Fig 1), it seems unlikely that variation capable of spanning the species differences is segregating in existing populations. On the other hand, given that pulse rate evolution is polygenic and effect sizes are small (e.g. this study), selective advantages of individual alleles should also be small. As such, standing genetic variation is a more likely source of beneficial, small-effect mutations because the short evolutionary timespan in which these species have diverged leaves little time for *de novo* mutations of this sort to arise (Hermisson and Pennings 2005; Barrett and Schluter 2008). Although parallel genetic mechanisms are more likely when standing genetic variation is involved (Barrett and Schluter 2008), both theoretical and empirical work on polygenic trait evolution more often suggest independent genetics (Elmer and Meyer 2011; Conte et al. 2012; Agrawal 2017). Thus, our results are an exciting counter example regardless of whether trait divergence is fueled by ancestral or *de novo* variants.

### Consolidating empirical observations with theory

Based on theory, we hypothesized that changes in common QTL between species pairs would be both limited and more likely for QTL of large effect (Orr 2005b; MacPherson and Nuismer 2017). In contrast, we found substantial QTL overlap that was unrelated to the magnitude of the QTL effect (Fig 2C). For instance, consider the shared QTL on LG5 that shows relatively large effect in KonPar (0.22 pps or 15.5% of the parental difference), as opposed to relatively small effect in CerEuk (0.07 pps or 4.90% of the parental difference). This suggests phenotypic effect sizes do not predict the likelihood of genetic parallelism in polygenic evolution, underscoring the assertion that parallel evolution of polygenic traits remains poorly understood.

The QTL on LG5 and other shared QTL that show varying effect sizes depending on the species involved also reveal important qualities of quantitative trait evolution. The QTL underlying differences in more distantly related species (that also have more divergent phenotypes), on average, show higher effect sizes than the orthologous QTL in more closely related species. Therefore, as we argued above, it is conceivable that QTL regions accumulate causal variants. Accumulation of variants across QTL within a single species pair may proceed haphazardly or, alternatively, the order may reflect the genetic line of least resistance along which phenotypic evolution is expected to occur (Schluter 1996). In both cases, there are several mechanisms that ultimately lead to a clustered genetic architecture, including genomic rearrangements (Yeaman 2013) and a competitive advantage of linked variants (that have relatively larger joint effects and thus higher fitness) over unlinked small-effect variants (Yeaman and Whitlock 2011). The QTL that have accumulated multiple, tightly linked, small effect variants will appear as if they are single variants of moderate to large effect. This may explain, in part, why larger effect loci are observed relatively frequently, while quantitative genetics theory predicts phenotypic evolution proceeds with small steps (i.e. adaptive walks) and can resolve the apparent contradiction between micro-mutational theory and ubiquitous observations of major effect loci.

## ACKNOWLEDGEMENTS

We are thankful to the members of the Shaw lab, in particular Mingzi Xu for her input on linkage mapping and QTL analysis. We also thank the Cornell Department of Neurobiology and Behavior, in particular Michael Sheehan and Sarah Miller for helpful discussion about this study. We further thank Karl Broman and the r/QTL Google group as well as Biostars and Stackoverflow communities for invaluable help in bioinformatics scripting and data analysis. We thank the National Science Foundation for funding. The authors declare no conflict of interest, financial or otherwise. This project was funded by the National Science Foundation, grant number IOS1257682 “The Genomic Architecture underlying Behavioral Isolation and Speciation’.

