## Supplemental Figure 1 for "Parallel genomic architecture underlies repeated sexual signal divergence in Hawaiian *Laupala* crickets"

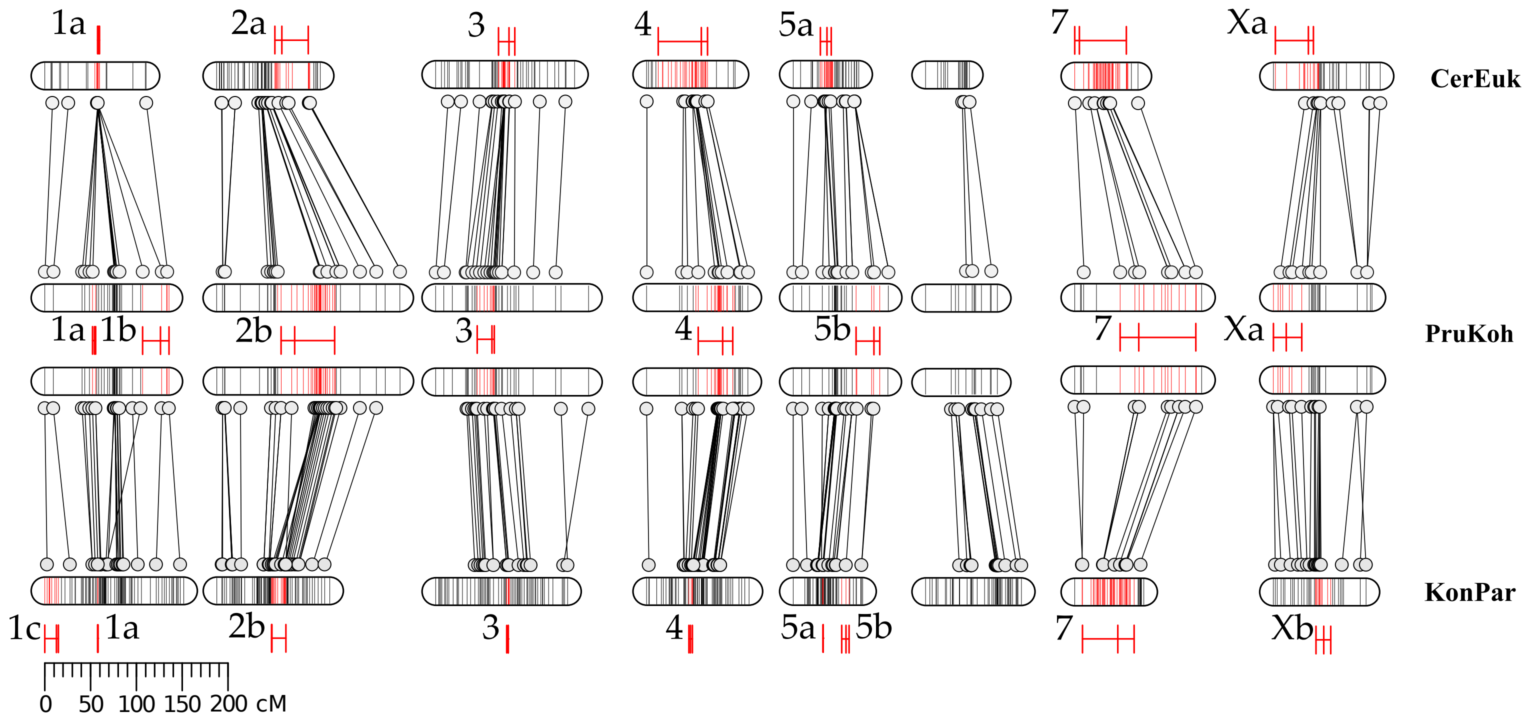


Fig S1. QTL overlap. The marker order and location of QTL are shown across the seven autosomal and X-linked linkage groups (LG) for all three crosses. The map for PruKoh is shown twice for comparative purposes. The scale bars give the scale of marker distances in cM. Markers (horizontal bars) are highlighted in red if they fall within the 95% Bayesian Credible Interval (red horizontal bars adjacent to the linkage groups). Homologous markers (from the same scaffold) between the crosses are connected by black lines.
