## Supplemental Figure 4 for "Parallel genomic architecture underlies repeated sexual signal divergence in Hawaiian *Laupala* crickets"

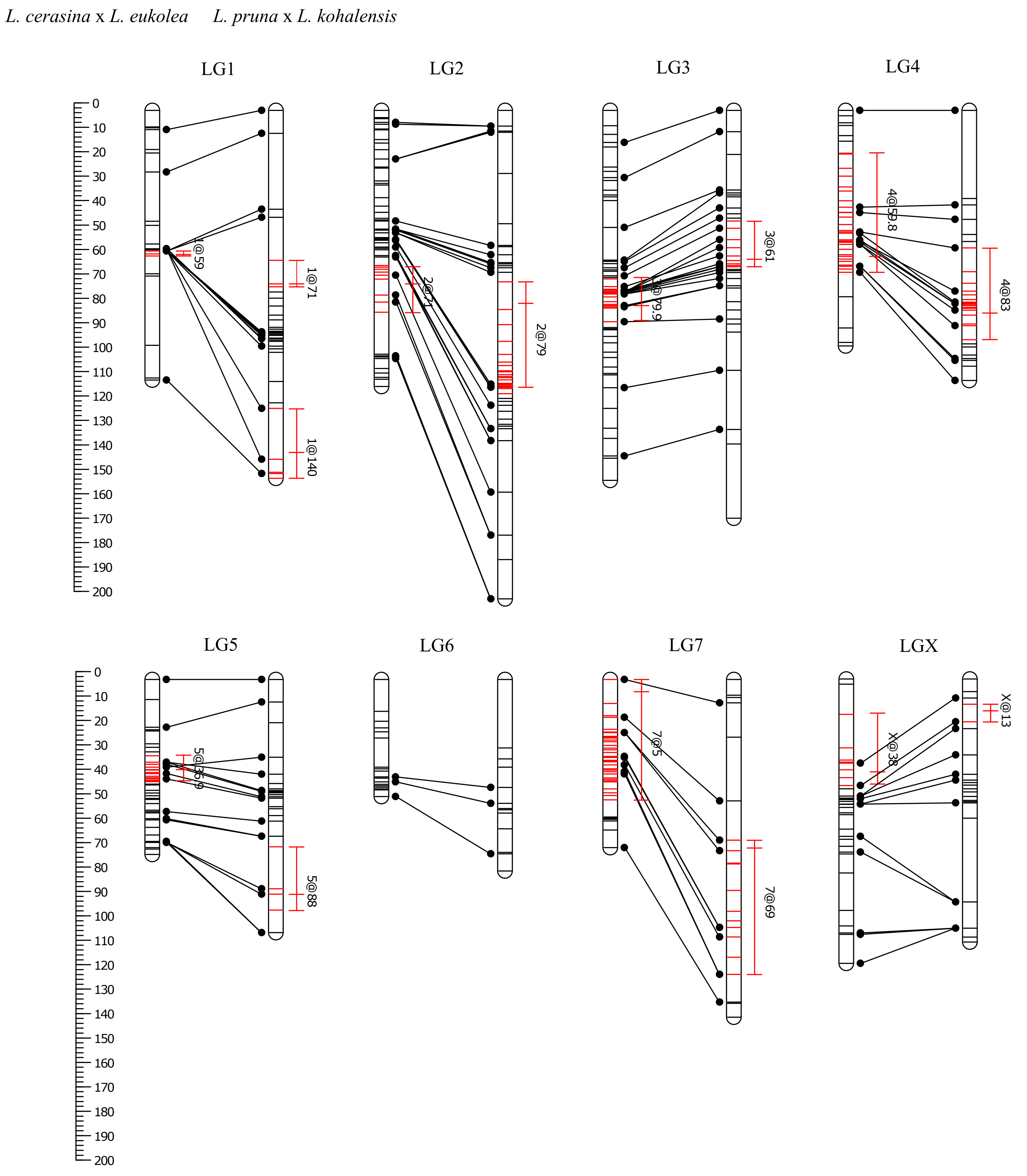


Fig S4. Linkage maps and QTL results for L. cerasina x L. eukolea and L. pruna x L. kohalensis. The marker order and location of QTL are shown across the seven autosomal and X-linked linkage groups (LG). The scale bars give the scale of marker distances in cM. Markers (horizontal bars) are highlighted in red if they fall within the 95% Bayesian Credible Interval (red horizontal bars adjacent to the linkage groups). Homologous markers (from the same scaffold) between the crosses are connected by black lines.
