## Supplementary Tables for "Parallel genomic architecture underlies repeated sexual signal divergence in Hawaiian *Laupala* crickets"

Table S1. QTL results for CerEuk. QTL were mapped using 230 F2 individuals. LG: linkage group. LOD: log-of-odds. A and B alleles denote *L. cerasina* and *L. eukolea* alleles, respectively. QTL effects are shown as the estimated effect in pulses per second derived from the multiple QTL model and as the percent variance explained relative to the parental difference (1.66 pps). All QTL effects are significantly different from zero (* P < 0.05; † P < 0.01).

|  |  |  |  |  | Genotypes | | |  |  |
| --- | --- | --- | --- | --- | --- | --- | --- | --- | --- |
| LG | position | LOD | nearest scaffold | Marker location | AA | AB | BB | effect (pps) | % of parental difference |
| 1 | 59.0 | 40.73 | S001131 | 58.8 | 2.89 | 3.17 | 3.31 | 0.203 | 12.23† |
| 2 | 71.0 | 4.78 | S001921 | 69.1 | 3.05 | 3.15 | 3.18 | 0.055 | 3.30† |
| 3 | 79.9 | 21.28 | S000385 | 79.9 | 3.02 | 3.14 | 3.23 | 0.123 | 7.40† |
| 4 | 59.8 | 3.97 | S016452 | 59.0 | 3.10 | 3.12 | 3.23 | 0.053 | 3.20† |
| 5 | 36.9 | 8.32 | S002445 | 36.9 | 3.05 | 3.16 | 3.19 | 0.068 | 4.08† |
| 7 | 5.0 | 3.51 | S007011 | 9.8 | 3.08 | 3.14 | 3.20 | 0.045 | 2.68† |
| X | 38.0 | 5.46 | S003132 | 37.2 | 3.10 | 3.17 | 3.10 | 0.041 | 2.46† |

Table S2. Scaffolds connected by red dashed lines in Fig 1D.

| QTL | shared scaffold | distance to the peak | | | QTL sharing |
| --- | --- | --- | --- | --- | --- |
|  |  | d(peak) CerEuk | d(peak) PruKoh | d(peak) KonPar |  |
| 1a | S002151 | 1.4 | 1.8 | 0 | QTL in all three |
| 1b | - | - | - | - | QTL unique to PruKoh |
| 1c | - | - | - | - | QTL unique to KonPar |
| 2a | - | - | - | - | QTL unique to CerEuk |
| 2b | S002839 | - | 6.3 | 4.3 | QTL not in CerEuk |
| 3 | S000529 | 4.5 | 2.5 | 0 | QTL in all three |
| 4 | S002870 | not on map | 3.5 | 2.4 | QTL in all three |
|  | S000182 | 5.8 | 2 | not on map | QTL in all three |
| 5a | S002565 | 2.4 | - | 0.6 | QTL not in PruKoh |
| 5b | S000770 | - | 17.3 | 0 | QTL not in CerEuk |
| 7 | S000677 | 16.8 | 4.2 | 15.6 | QTL in all three |
| Xa | S003455 | 5.6 | 3.5 | - | QTL not in KonPar |
| Xb | - | - | - | - | QTL unique to KonPar |

Table S3. 95% Bayesian Credible Intervals around the QTL peaks for all three crosses.

| QTL | position | scaffold | cross | comment |
| --- | --- | --- | --- | --- |
| 1a | 57.60005 | S004771 | cereuk |  |
| 1a | 57.60005 | S002151 | cereuk | shared with both |
| 1a | 57.60005 | S002242 | cereuk |  |
| 1a | 57.60005 | S001056 | cereuk |  |
| 1a | 57.60005 | S003828 | cereuk |  |
| 1a | 57.60006 | S000277 | cereuk |  |
| 1a | 57.60006 | S004794 | cereuk |  |
| 1a | 57.60006 | S002553 | cereuk |  |
| 1a | 57.60006 | S001522 | cereuk |  |
| 1a | 57.60006 | S001990 | cereuk |  |
| 1a | 57.60006 | S000828 | cereuk |  |
| 1a | 57.60006 | S000658 | cereuk |  |
| 1a | 57.60006 | S002761 | cereuk |  |
| 1a | 57.60006 | S003866 | cereuk |  |
| 1a | 57.60006 | S003095 | cereuk |  |
| 1a | 57.60007 | S008331 | cereuk |  |
| 1a | 57.60007 | S000409 | cereuk |  |
| 1a | 57.60007 | S002490 | cereuk |  |
| 1a | 57.60007 | S001589 | cereuk |  |
| 1a | 57.60007 | S000270 | cereuk |  |
| 1a | 57.60007 | S001131 | cereuk |  |
| 1a | 58.80007 | S001131 | cereuk |  |
| 1a | 59.60007 | S005462 | cereuk | peak |
| 1a | 57.60003 | S003640 | konpar |  |
| 1a | 58.10003 | S002151 | konpar | peak/shared with both |
| 1a | 48.5 | S001056 | prukoh |  |
| 1a | 52.20001 | S002151 | prukoh | shared with both |
| 1a | 55.40001 | S000873 | prukoh | peak |
| 1b | 106.7 | S002518 | prukoh |  |
| 1b | 127.5 | S002077 | prukoh | peak |
| 1b | 132.9 | S004757 | prukoh |  |
| 1b | 133.4 | S001067 | prukoh |  |
| 1b | 135.3 | S014985 | prukoh |  |
| 1c | 0 | S021409 | konpar |  |
| 1c | 2.600001 | S003488 | konpar |  |
| 1c | 2.600002 | S002234 | konpar |  |
| 1c | 4.500003 | S000390 | konpar |  |
| 1c | 4.900004 | S000390 | konpar |  |
| 1c | 5.300005 | S000390 | konpar |  |
| 1c | 7.800006 | S005944 | konpar |  |
| 1c | 11.00001 | S000584 | konpar |  |
| 1c | 12.70001 | S002179 | konpar | peak |
| 1c | 14.70001 | S000397 | konpar |  |
| 2a | 63.50006 | S010389 | cereuk |  |
| 2a | 64.20007 | S001526 | cereuk |  |
| 2a | 65.00007 | S001272 | cereuk |  |
| 2a | 66.20007 | S004513 | cereuk |  |
| 2a | 67.50007 | S000482 | cereuk |  |
| 2a | 67.50007 | S001921 | cereuk |  |
| 2a | 69.10007 | S001921 | cereuk | peak |
| 2a | 75.60007 | S001602 | cereuk |  |
| 2a | 78.50007 | S001602 | cereuk |  |
| 2a | 82.60007 | S000793 | cereuk |  |
| 2a | 82.60007 | S000793 | cereuk |  |
| 2a | 99.80008 | S009494 | cereuk |  |
| 2a | 100.5001 | S000326 | cereuk |  |
| 2b | 58.70005 | S002456 | konpar |  |
| 2b | 59.70005 | S005289 | konpar |  |
| 2b | 59.70005 | S000230 | konpar |  |
| 2b | 60.10006 | S000517 | konpar | peak |
| 2b | 60.70006 | S004070 | konpar |  |
| 2b | 61.30006 | S002169 | konpar |  |
| 2b | 61.90006 | S003079 | konpar |  |
| 2b | 63.20006 | S003079 | konpar |  |
| 2b | 64.30006 | S002839 | konpar | shared with prukoh |
| 2b | 66.40006 | S001230 | konpar |  |
| 2b | 70.10006 | S003202 | konpar |  |
| 2b | 72.00006 | S000819 | konpar |  |
| 2b | 72.70006 | S000209 | konpar |  |
| 2b | 73.50007 | S000588 | konpar |  |
| 2b | 73.50007 | S004807 | konpar |  |
| 2b | 73.50007 | S003032 | konpar |  |
| 2b | 74.10007 | S001171 | konpar |  |
| 2b | 74.10007 | S002243 | konpar |  |
| 2b | 74.50007 | S001327 | konpar |  |
| 2b | 74.50007 | S000330 | konpar |  |
| 2b | 74.70007 | S000883 | konpar |  |
| 2b | 74.70007 | S002156 | konpar |  |
| 2b | 75.40007 | S000991 | konpar |  |
| 2b | 46.40001 | S001227 | prukoh |  |
| 2b | 55.30001 | S000615 | prukoh |  |
| 2b | 55.30001 | S001126 | prukoh |  |
| 2b | 55.80001 | S001192 | prukoh |  |
| 2b | 59.10001 | S000416 | prukoh |  |
| 2b | 62.30001 | S001514 | prukoh |  |
| 2b | 62.80001 | S007459 | prukoh |  |
| 2b | 63.60001 | S002088 | prukoh |  |
| 2b | 64.40001 | S002097 | prukoh |  |
| 2b | 66.30001 | S001659 | prukoh |  |
| 2b | 70.20002 | S002376 | prukoh |  |
| 2b | 81.50002 | S000991 | prukoh |  |
| 2b | 87.70002 | S001378 | prukoh |  |
| 2b | 94.60002 | S006692 | prukoh |  |
| 2b | 99.90002 | S004212 | prukoh |  |
| 2b | 103 | S002877 | prukoh |  |
| 2b | 104.5 | S001532 | prukoh | peak |
| 2b | 106.6 | S001171 | prukoh |  |
| 2b | 107 | S003735 | prukoh |  |
| 2b | 108.2 | S003735 | prukoh |  |
| 2b | 108.2 | S002243 | prukoh |  |
| 2b | 109 | S000302 | prukoh |  |
| 2b | 109.4 | S003094 | prukoh |  |
| 2b | 110.3 | S002839 | prukoh | shared with konpar |
| 2b | 111.8 | S000434 | prukoh |  |
| 2b | 112.1 | S000517 | prukoh |  |
| 2b | 112.3 | S004769 | prukoh |  |
| 2b | 112.3 | S000624 | prukoh |  |
| 2b | 112.8 | S003079 | prukoh |  |
| 2b | 112.8 | S004663 | prukoh |  |
| 2b | 113.1 | S010869 | prukoh |  |
| 2b | 113.1 | S010879 | prukoh |  |
| 2b | 113.1 | S003182 | prukoh |  |
| 2b | 113.1 | S003202 | prukoh |  |
| 2b | 113.1 | S000279 | prukoh |  |
| 2b | 113.1 | S003118 | prukoh |  |
| 2b | 113.1 | S003118 | prukoh |  |
| 2b | 113.4 | S000330 | prukoh |  |
| 2b | 113.9 | S000883 | prukoh |  |
| 2b | 116 | S000842 | prukoh |  |
| 2b | 117.9 | S001629 | prukoh |  |
| 2b | 119.1 | S002295 | prukoh |  |
| 2b | 120.7 | S013864 | prukoh |  |
| 2b | 123.2 | S000199 | prukoh |  |
| 2b | 126.4 | S001797 | prukoh |  |
| 2b | 128.5001 | S001483 | prukoh |  |
| 3 | 68.40003 | S001400 | cereuk |  |
| 3 | 69.00003 | S004777 | cereuk |  |
| 3 | 72.10003 | S002774 | cereuk |  |
| 3 | 73.20003 | S002002 | cereuk |  |
| 3 | 73.70003 | S002500 | cereuk |  |
| 3 | 74.20004 | S002566 | cereuk |  |
| 3 | 74.20004 | S000749 | cereuk |  |
| 3 | 74.40004 | S000558 | cereuk |  |
| 3 | 74.40004 | S000236 | cereuk |  |
| 3 | 74.40004 | S000454 | cereuk |  |
| 3 | 74.40004 | S001309 | cereuk |  |
| 3 | 74.40004 | S001309 | cereuk |  |
| 3 | 74.40004 | S001148 | cereuk |  |
| 3 | 74.40004 | S002598 | cereuk |  |
| 3 | 74.40004 | S000500 | cereuk |  |
| 3 | 74.40005 | S003066 | cereuk |  |
| 3 | 74.40005 | S000529 | cereuk | shared with both |
| 3 | 74.60005 | S006750 | cereuk |  |
| 3 | 74.60005 | S001177 | cereuk |  |
| 3 | 74.60005 | S001652 | cereuk |  |
| 3 | 74.60005 | S003635 | cereuk |  |
| 3 | 74.60005 | S003635 | cereuk |  |
| 3 | 74.60005 | S000917 | cereuk |  |
| 3 | 74.60005 | S004098 | cereuk |  |
| 3 | 74.60005 | S002879 | cereuk |  |
| 3 | 74.60006 | S002027 | cereuk |  |
| 3 | 74.80006 | S005557 | cereuk |  |
| 3 | 74.80006 | S003170 | cereuk |  |
| 3 | 75.20006 | S003534 | cereuk |  |
| 3 | 75.20006 | S000212 | cereuk |  |
| 3 | 75.20006 | S000212 | cereuk |  |
| 3 | 75.20006 | S000550 | cereuk |  |
| 3 | 75.20006 | S001331 | cereuk |  |
| 3 | 75.20006 | S005448 | cereuk |  |
| 3 | 75.20006 | S006775 | cereuk |  |
| 3 | 76.40007 | S001260 | cereuk |  |
| 3 | 78.30007 | S000656 | cereuk |  |
| 3 | 78.30007 | S003481 | cereuk |  |
| 3 | 79.40007 | S001265 | cereuk |  |
| 3 | 79.90007 | S000835 | cereuk | peak |
| 3 | 79.90007 | S000385 | cereuk |  |
| 3 | 80.40007 | S000385 | cereuk |  |
| 3 | 80.40007 | S003125 | cereuk |  |
| 3 | 80.90007 | S002439 | cereuk |  |
| 3 | 86.50007 | S000203 | cereuk |  |
| 3 | 77.60005 | S002665 | konpar |  |
| 3 | 79.10005 | S001173 | konpar |  |
| 3 | 79.10005 | S002566 | konpar |  |
| 3 | 79.10005 | S005998 | konpar | peak |
| 3 | 79.10005 | S000529 | konpar | shared with both |
| 3 | 45.30001 | S001130 | prukoh |  |
| 3 | 48.30001 | S000927 | prukoh |  |
| 3 | 52.90001 | S000558 | prukoh |  |
| 3 | 56.20001 | S000558 | prukoh |  |
| 3 | 59.60001 | S002774 | prukoh |  |
| 3 | 61.50002 | S000726 | prukoh | peak |
| 3 | 62.80002 | S000749 | prukoh |  |
| 3 | 62.80002 | S000361 | prukoh |  |
| 3 | 63.50002 | S002851 | prukoh |  |
| 3 | 64.00002 | S000529 | prukoh | shared with both |
| 3 | 64.00002 | S002500 | prukoh |  |
| 4 | 12.60001 | S000309 | cereuk |  |
| 4 | 17.50001 | S005469 | cereuk |  |
| 4 | 17.90001 | S005469 | cereuk |  |
| 4 | 23.70001 | S001747 | cereuk |  |
| 4 | 26.90001 | S003830 | cereuk |  |
| 4 | 26.90001 | S002318 | cereuk |  |
| 4 | 31.30001 | S000426 | cereuk |  |
| 4 | 33.00001 | S013482 | cereuk |  |
| 4 | 37.00001 | S002317 | cereuk |  |
| 4 | 39.60001 | S000376 | cereuk |  |
| 4 | 41.70002 | S003329 | cereuk |  |
| 4 | 41.80002 | S001151 | cereuk |  |
| 4 | 43.70002 | S000496 | cereuk |  |
| 4 | 46.70002 | S003881 | cereuk |  |
| 4 | 49.20002 | S005538 | cereuk |  |
| 4 | 49.80002 | S001965 | cereuk |  |
| 4 | 50.30002 | S000668 | cereuk |  |
| 4 | 53.00002 | S011082 | cereuk |  |
| 4 | 53.10002 | S006899 | cereuk |  |
| 4 | 53.10002 | S000802 | cereuk |  |
| 4 | 53.10003 | S014891 | cereuk |  |
| 4 | 53.10003 | S000900 | cereuk |  |
| 4 | 53.10003 | S001783 | cereuk |  |
| 4 | 53.10003 | S001282 | cereuk |  |
| 4 | 53.60003 | S004811 | cereuk |  |
| 4 | 53.60003 | S000836 | cereuk |  |
| 4 | 54.00003 | S002061 | cereuk |  |
| 4 | 54.00003 | S000182 | cereuk | shared with prukoh |
| 4 | 54.90003 | S001754 | cereuk |  |
| 4 | 54.90003 | S010013 | cereuk |  |
| 4 | 56.90004 | S001243 | cereuk |  |
| 4 | 59.10004 | S002058 | cereuk |  |
| 4 | 59.80004 | S016452 | cereuk | peak |
| 4 | 60.90004 | S000591 | cereuk |  |
| 4 | 60.90004 | S000455 | cereuk |  |
| 4 | 61.80004 | S001530 | cereuk |  |
| 4 | 63.30004 | S000422 | cereuk |  |
| 4 | 63.80004 | S000674 | cereuk |  |
| 4 | 64.90004 | S000999 | cereuk |  |
| 4 | 66.30004 | S001412 | cereuk |  |
| 4 | 46.20002 | S002556 | konpar |  |
| 4 | 48.80002 | S001965 | konpar | peak |
| 4 | 48.80003 | S001614 | konpar |  |
| 4 | 50.00003 | S000730 | konpar |  |
| 4 | 50.00003 | S010617 | konpar |  |
| 4 | 50.40003 | S002870 | konpar | shared with prukoh |
| 4 | 50.8 | S001617 | prukoh |  |
| 4 | 53.70001 | S002556 | prukoh |  |
| 4 | 56.30001 | S001965 | prukoh |  |
| 4 | 66.00001 | S000301 | prukoh |  |
| 4 | 70.80001 | S007730 | prukoh |  |
| 4 | 74.00001 | S000836 | prukoh |  |
| 4 | 75.50001 | S014123 | prukoh |  |
| 4 | 77.50001 | S000730 | prukoh |  |
| 4 | 77.50001 | S002870 | prukoh | shared with konpar |
| 4 | 78.50001 | S000668 | prukoh |  |
| 4 | 78.80001 | S006241 | prukoh |  |
| 4 | 79.10002 | S001482 | prukoh |  |
| 4 | 79.10002 | S001401 | prukoh |  |
| 4 | 79.10002 | S003206 | prukoh |  |
| 4 | 79.10002 | S001416 | prukoh |  |
| 4 | 79.10002 | S000490 | prukoh |  |
| 4 | 79.10002 | S006899 | prukoh |  |
| 4 | 79.10002 | S001634 | prukoh |  |
| 4 | 79.10002 | S000802 | prukoh |  |
| 4 | 79.10002 | S001005 | prukoh |  |
| 4 | 79.10002 | S000182 | prukoh | shared with cereuk |
| 4 | 79.10003 | S001609 | prukoh |  |
| 4 | 79.10003 | S001860 | prukoh |  |
| 4 | 79.10003 | S014891 | prukoh |  |
| 4 | 79.40003 | S001277 | prukoh |  |
| 4 | 79.40003 | S002038 | prukoh |  |
| 4 | 79.90003 | S006623 | prukoh |  |
| 4 | 80.70003 | S002090 | prukoh |  |
| 4 | 81.00003 | S002090 | prukoh | peak |
| 4 | 81.00003 | S009434 | prukoh |  |
| 4 | 81.00003 | S009434 | prukoh |  |
| 4 | 81.80004 | S001754 | prukoh |  |
| 4 | 83.80004 | S003111 | prukoh |  |
| 4 | 87.50004 | S009873 | prukoh |  |
| 5a | 29.60001 | S002808 | cereuk |  |
| 5a | 31.20001 | S000562 | cereuk |  |
| 5a | 31.20001 | S002687 | cereuk |  |
| 5a | 33.90001 | S001294 | cereuk |  |
| 5a | 33.90001 | S000366 | cereuk |  |
| 5a | 33.90002 | S000430 | cereuk |  |
| 5a | 34.70002 | S002617 | cereuk |  |
| 5a | 34.70002 | S003777 | cereuk |  |
| 5a | 34.70002 | S000359 | cereuk |  |
| 5a | 34.70002 | S004947 | cereuk |  |
| 5a | 35.90002 | S000745 | cereuk |  |
| 5a | 35.90002 | S001510 | cereuk |  |
| 5a | 36.90002 | S002445 | cereuk | peak |
| 5a | 37.10002 | S001239 | cereuk |  |
| 5a | 38.10002 | S001239 | cereuk |  |
| 5a | 38.10003 | S000272 | cereuk |  |
| 5a | 38.50003 | S002565 | cereuk | shared with konpar |
| 5a | 39.60003 | S001371 | cereuk |  |
| 5a | 40.20003 | S001371 | cereuk |  |
| 5a | 40.70003 | S003497 | cereuk |  |
| 5a | 40.70003 | S001044 | cereuk |  |
| 5a | 41.30003 | S000933 | cereuk |  |
| 5a | 41.50003 | S000933 | cereuk |  |
| 5a | 32.00003 | S002944 | konpar |  |
| 5a | 32.10003 | S000820 | konpar |  |
| 5a | 32.50003 | S004383 | konpar | peak |
| 5a | 32.80003 | S004683 | konpar |  |
| 5a | 33.10003 | S002565 | konpar | shared with cereuk |
| 5b | 53.10006 | S001803 | konpar |  |
| 5b | 57.60006 | S000770 | konpar | peak / shared with prukoh |
| 5b | 61.10006 | S004681 | konpar |  |
| 5b | 69.60003 | S000770 | prukoh | shared with konpar |
| 5b | 85.70003 | S007270 | prukoh |  |
| 5b | 87.80003 | S007270 | prukoh | peak |
| 5b | 94.30003 | S001100 | prukoh |  |
| 5b | 103.5 | S003304 | prukoh |  |
| 7 | 0 | S000257 | cereuk |  |
| 7 | 1.00E-06 | S003103 | cereuk |  |
| 7 | 9.800002 | S007011 | cereuk | peak |
| 7 | 14.8 | S000628 | cereuk |  |
| 7 | 15.5 | S000628 | cereuk |  |
| 7 | 15.50001 | S000633 | cereuk |  |
| 7 | 20.40001 | S003151 | cereuk |  |
| 7 | 21.50001 | S003759 | cereuk |  |
| 7 | 21.80001 | S000677 | cereuk | shared with both |
| 7 | 23.30001 | S005499 | cereuk |  |
| 7 | 23.60001 | S001317 | cereuk |  |
| 7 | 24.00001 | S000183 | cereuk |  |
| 7 | 24.80001 | S000294 | cereuk |  |
| 7 | 25.50001 | S003115 | cereuk |  |
| 7 | 27.00001 | S003115 | cereuk |  |
| 7 | 28.10002 | S000195 | cereuk |  |
| 7 | 28.80002 | S003292 | cereuk |  |
| 7 | 29.90002 | S000712 | cereuk |  |
| 7 | 29.90002 | S004067 | cereuk |  |
| 7 | 31.50002 | S004319 | cereuk |  |
| 7 | 31.50002 | S007241 | cereuk |  |
| 7 | 31.50002 | S001018 | cereuk |  |
| 7 | 31.90002 | S001018 | cereuk |  |
| 7 | 31.90002 | S001018 | cereuk |  |
| 7 | 33.10002 | S003971 | cereuk |  |
| 7 | 33.40003 | S002652 | cereuk |  |
| 7 | 33.40003 | S002652 | cereuk |  |
| 7 | 33.40003 | S002652 | cereuk |  |
| 7 | 33.70003 | S002087 | cereuk |  |
| 7 | 35.00003 | S002441 | cereuk |  |
| 7 | 36.60003 | S001865 | cereuk |  |
| 7 | 37.10003 | S001612 | cereuk |  |
| 7 | 37.80003 | S004540 | cereuk |  |
| 7 | 37.80003 | S004540 | cereuk |  |
| 7 | 37.80003 | S000442 | cereuk |  |
| 7 | 37.80004 | S004909 | cereuk |  |
| 7 | 38.70004 | S004909 | cereuk |  |
| 7 | 40.30004 | S001259 | cereuk |  |
| 7 | 40.30004 | S001600 | cereuk |  |
| 7 | 40.30004 | S003542 | cereuk |  |
| 7 | 40.90004 | S004220 | cereuk |  |
| 7 | 40.90004 | S010292 | cereuk |  |
| 7 | 41.50004 | S000610 | cereuk |  |
| 7 | 41.90004 | S004936 | cereuk |  |
| 7 | 44.80004 | S002507 | cereuk |  |
| 7 | 46.40005 | S003682 | cereuk |  |
| 7 | 47.50005 | S001645 | cereuk |  |
| 7 | 49.30005 | S001744 | cereuk |  |
| 7 | 56.30005 | S001482 | cereuk |  |
| 7 | 8.400002 | S003249 | konpar |  |
| 7 | 17.9 | S002304 | konpar |  |
| 7 | 20.5 | S002736 | konpar |  |
| 7 | 21.30001 | S002736 | konpar |  |
| 7 | 23.30001 | S002217 | konpar |  |
| 7 | 24.40001 | S000628 | konpar |  |
| 7 | 25.20001 | S000628 | konpar |  |
| 7 | 25.30001 | S000628 | konpar |  |
| 7 | 26.00001 | S003860 | konpar |  |
| 7 | 27.60001 | S002592 | konpar |  |
| 7 | 28.10001 | S002165 | konpar |  |
| 7 | 30.00001 | S001362 | konpar |  |
| 7 | 30.90001 | S000677 | konpar | shared with both |
| 7 | 31.20002 | S000677 | konpar |  |
| 7 | 31.20002 | S000677 | konpar |  |
| 7 | 32.00002 | S001317 | konpar |  |
| 7 | 32.20002 | S001317 | konpar |  |
| 7 | 33.20002 | S006047 | konpar |  |
| 7 | 35.70002 | S003521 | konpar |  |
| 7 | 38.70002 | S003115 | konpar |  |
| 7 | 39.30002 | S002926 | konpar |  |
| 7 | 39.50002 | S001456 | konpar |  |
| 7 | 39.80002 | S003292 | konpar |  |
| 7 | 40.00003 | S000195 | konpar |  |
| 7 | 40.70003 | S001907 | konpar |  |
| 7 | 41.80003 | S001701 | konpar |  |
| 7 | 42.30003 | S000712 | konpar |  |
| 7 | 42.70003 | S000450 | konpar |  |
| 7 | 42.70003 | S000450 | konpar |  |
| 7 | 43.50003 | S001018 | konpar |  |
| 7 | 46.30003 | S001185 | konpar | peak |
| 7 | 48.70003 | S002087 | konpar |  |
| 7 | 49.50003 | S003075 | konpar |  |
| 7 | 49.50004 | S008198 | konpar |  |
| 7 | 49.70004 | S002441 | konpar |  |
| 7 | 50.50004 | S001071 | konpar |  |
| 7 | 51.10004 | S004702 | konpar |  |
| 7 | 51.10004 | S004853 | konpar |  |
| 7 | 52.70004 | S003063 | konpar |  |
| 7 | 54.10004 | S000137 | konpar |  |
| 7 | 55.50004 | S002309 | konpar |  |
| 7 | 55.90004 | S002309 | konpar |  |
| 7 | 56.50004 | S000442 | konpar |  |
| 7 | 56.70005 | S004540 | konpar |  |
| 7 | 56.80005 | S004823 | konpar |  |
| 7 | 56.90005 | S004909 | konpar |  |
| 7 | 58.00005 | S001259 | konpar |  |
| 7 | 59.90005 | S004976 | konpar |  |
| 7 | 60.50005 | S004935 | konpar |  |
| 7 | 64.70005 | S002507 | konpar |  |
| 7 | 64.70005 | S004334 | konpar |  |
| 7 | 23.6 | S004384 | prukoh |  |
| 7 | 49.70001 | S000633 | prukoh |  |
| 7 | 65.80001 | S000677 | prukoh | shared with both |
| 7 | 70.10001 | S000677 | prukoh | peak |
| 7 | 70.10001 | S000677 | prukoh |  |
| 7 | 75.20001 | S001606 | prukoh |  |
| 7 | 75.50001 | S001606 | prukoh |  |
| 7 | 86.40001 | S001531 | prukoh |  |
| 7 | 94.90001 | S003448 | prukoh |  |
| 7 | 98.80001 | S000338 | prukoh |  |
| 7 | 101.5 | S001018 | prukoh |  |
| 7 | 101.5 | S001018 | prukoh |  |
| 7 | 105.4 | S002441 | prukoh |  |
| 7 | 113.7 | S002309 | prukoh |  |
| 7 | 120.7 | S004909 | prukoh |  |
| 7 | 132.1 | S000429 | prukoh |  |
| Xa | 2.100003 | S004528 | cereuk |  |
| Xa | 14.5 | S000936 | cereuk |  |
| Xa | 28.20001 | S000108 | cereuk |  |
| Xa | 33.20001 | S000913 | cereuk |  |
| Xa | 34.00001 | S000188 | cereuk |  |
| Xa | 34.50001 | S001780 | cereuk |  |
| Xa | 37.20001 | S003132 | cereuk |  |
| Xa | 37.20001 | S003132 | cereuk | peak |
| Xa | 40.40001 | S001725 | cereuk |  |
| Xa | 43.60001 | S003455 | cereuk | shared with prukoh |
| Xa | 7.800005 | S001780 | prukoh |  |
| Xa | 10.30001 | S001105 | prukoh |  |
| Xa | 17.50001 | S003455 | prukoh | peak / shared with cereuk |
| Xa | 20.30001 | S001183 | prukoh |  |
| Xa | 31.10001 | S000219 | prukoh |  |
| Xb | 46.60003 | S003053 | konpar |  |
| Xb | 46.60003 | S001257 | konpar |  |
| Xb | 46.60003 | S000441 | konpar |  |
| Xb | 46.60003 | S000613 | konpar |  |
| Xb | 46.60003 | S000613 | konpar |  |
| Xb | 47.00003 | S005739 | konpar |  |
| Xb | 47.00004 | S002517 | konpar |  |
| Xb | 47.00004 | S003732 | konpar |  |
| Xb | 47.00004 | S000364 | konpar |  |
| Xb | 47.00004 | S000275 | konpar |  |
| Xb | 47.00004 | S000275 | konpar |  |
| Xb | 47.00004 | S003307 | konpar |  |
| Xb | 48.50004 | S002191 | konpar |  |
| Xb | 50.10004 | S001930 | konpar |  |
| Xb | 50.40004 | S001930 | konpar |  |
| Xb | 50.40004 | S001930 | konpar |  |
| Xb | 50.80005 | S001930 | konpar |  |
| Xb | 50.80005 | S002003 | konpar |  |
| Xb | 51.90005 | S004832 | konpar |  |
| Xb | 51.90005 | S005630 | konpar |  |
| Xb | 53.40005 | S004078 | konpar |  |
| Xb | 53.40005 | S001055 | konpar |  |
| Xb | 59.20005 | S001995 | konpar |  |
| Xb | 62.60005 | S001995 | konpar |  |

Table S4. Extent of QTL sharing. The number and location of unique QTL to the KonPar, PruKoh, and CerEuk crosses, the number and location of double QTL (present in two out of three crosses) and the number and location of triple QTL are shown. See Fig S2-S4 for overlap in BCI.

|  | LG | position (cM) in KonPar | position (cM) in PruKoh | position (cM) in CerEuk |
| --- | --- | --- | --- | --- |
| unique | 1 | - | 127.5 | - |
|  | 1 | 12.7 | - | - |
|  | 2 | - | - | 71 |
|  | X | 55 | - | - |
| double | 2 | 60 | 104 |  |
|  | 5 | 57.6 | 57 | - |
|  | 5 | 32.5 | - | 36.9 |
|  | X | - | 14 | 38 |
| triple | 1 | 58.1 | 54 | 59 |
|  | 3 | 79.1 | 61.5 | 79.9 |
|  | 4 | 48 | 81 | 59.8 |
